# Cellular moonlighting function of Hsp20 directs morphological and pathogenic development in *Ustilago maydis*

**DOI:** 10.1101/2024.11.08.622622

**Authors:** Aroni Mitra, Ankita Kar, Koustav Bhakta, Anisha Roy, Dibya Mukherjee, Abhrajyoti Ghosh, Anupama Ghosh

## Abstract

*Ustilago maydis* Hsp20 is involved in the pathogenicity of the fungus. In this study we have investigated the molecular basis of contribution of Hsp20 to *U. maydis* pathogenicity. Through biochemical studies we have demonstrated environment-dependent oligomeric plasticity associated with Hsp20. Hsp20 was also found to form higher order oligomers that undergo phase separation in vitro. Within cells Hsp20 was found to form distinct punctate structures that we believe play a pivotal role in its function. These punctate structures were demonstrated to sequester proteins such as actin and septin within it. Absence of Hsp20 was found to significantly affect key cellular processes like endocytosis, budding, cell polarity determination and mating in *U. maydis* cells. The deletion mutant failed to sporulate and complete pathogenic life cycle. This study presents a comprehensive understanding of the pathogenic development of *U. maydis* in reference to the moonlighting function of Hsp20 within the cell.

## Introduction

Small heat shock proteins (sHsp) are ATP-independent chaperones (Jakob *et al*., 1993) that exhibit environment-regulated oligomeric plasticity (Stengel *et al*., 2010, Roy *et al*., 2018). Usually, these proteins are stored within the cell as higher order oligomers. Under specific conditions leading to cellular stress, functionally active dimeric forms of the protein are produced. sHsps, therefore, serve primarily as stress-responsive proteins that relieve a stressed cell from the load of unfolded or misfolded functionally inactive proteins. Therefore, the major cellular function influenced by a particular sHsp depends on its client proteins. For instance, sHsps are responsive to mechanical stress to the cell owing to their interaction with different cytoskeletal elements as client proteins. Examples include association of HspB1 with keratin that exhibits extensive remodelling in response to mechanical stress (Kayser *et al*., 2013). In addition, HspB1 also exhibits interaction with an actin-binding protein filamin, involved in musculosketal mechanosensing (Collier & Benesch, 2020). In mammals sHsps inhibit apoptosis by targeting several proteins that regulate critical events in the underlying molecular signalling pathway. HspB1, for instance, binds and sequesters cytochrome c, making it unavailable for activation of Apaf1 to form apoptosomes. Apoptosomes in turn initiate the activation of caspases leading to execution of the apoptotic process (Bruey *et al*., 2000). A recent study revealed yet another function of sHsp in the cellular endocytosis process. In this study, HspA8 was found to be involved in the attachment of porcine reproductive and respiratory syndrome virus (PRRSV) to the host cell and its subsequent internalization through a clathrin dependent endocytosis pathway (Wang *et al*., 2022). Besides proteins, sHsps have also been demonstrated to exhibit interaction with lipid components of the membranes, leading to membrane stabilizing functions. For instance, both bacterial sHsp Lo18 and mammalian HspB1 exhibit interaction with phospholipids and maintain membrane integrity under conditions of heat stress (Coucheney *et al*., 2005, Csoboz *et al*., 2022). Similarly, an intrinsically disordered stress protein Hsp12 from *Saccharomyces cerevisiae* and *Ustilago maydis* have been shown to modulate membrane functions through their interaction with the membrane lipids (Welker *et al*., 2010, Mitra *et al*., 2023). Thus, sHsps provide protection to a host cell by bringing stability to major cellular processes under conditions of stress. In this study we present experimental evidences indicating key role of *U. maydis* Hsp20 in a number of cellular processes in *U. maydis* including mating, endocytosis, maintaining actin cytoskeletal dynamics and determining cell polarity during budding. Together, all of these cellular functions defined the involvement Hsp20 in the morphological and pathogenic development of *U. maydis*. The detailed molecular mechanism explaining the contribution of Hsp20 in these major cellular processes however, remains elusive. Nevertheless, an exhaustive study of the biochemical properties of purified *U. maydis* Hsp20 revealed a protein aggregation prevention activity associated with the protein. In addition, Hsp20 was also demonstrated to exhibit temperature and hydrophobic microenvironment induced oligomeric plasticity. Under conditions of induced molecular crowding in vitro, purified Hsp20 also showed formation of liquid condensates that can sequester substrate protein within it. Taken together these data provide important cues for further studies relating the molecular mechanism of action of Hsp20 with its biological function in *U. maydis*.

## Results

### *Ustilago maydis* Hsp20 is a conventional small heat shock protein with a conserved alpha crystalline domain

*U. maydis* Hsp20 is a 215-residue small heat shock protein (sHsp) with an alpha crystalline/Hsp20 domain (Interpro IPR002068) spanning from D48 to T214 (Fig 1A). Previous studies have demonstrated that sHsps form higher-order oligomers as storage forms of the protein. Under proteotoxic stress, such as heat shock or exposure to a hydrophobic environment, the protein releases dimeric or tetrameric forms that prevent protein aggregation (Roy et al., 2018, Santhanagopalan *et al*., 2018). We aimed to test whether *U. maydis* Hsp20 also exhibits oligomerization dynamics in response to environmental conditions. Recombinant His_6X_-Hsp20 was purified to homogeneity from *E. coli* BL21 DE3 cells (Fig S1). The protein was analyzed using dynamic light scattering (DLS) to examine its oligomeric plasticity. At room temperature, the protein formed large oligomers with a hydrodynamic diameter of 13.54 nm (Fig 1B). When the temperature was increased to mimic heat shock conditions, the hydrodynamic diameter gradually decreased, suggesting a transition from higher-order oligomers to lower-order ones. Previous studies also identified a C-terminal IXI/V motif responsible for higher-order oligomer formation in sHsps (Studer *et al*., 2002, Saji *et al*., 2007). Upon analyzing the *U. maydis* Hsp20 sequence, we identified an IAI motif within residues 209 to 211. To investigate the role of this motif in oligomerization, it was mutated to KKK, and DLS was performed. The His_6X_-Hsp20_KKK_ mutant formed smaller oligomers at all temperatures (Fig 1B), indicating that the IAI motif is crucial for higher-order oligomer formation. In addition to temperature, we sought to mimic the hydrophobic microenvironment created during proteotoxic stress. So, the protein was treated with 0.4 mM SDS, and DLS revealed that SDS exposure led to the formation of smaller oligomers with a lower hydrodynamic diameter (Fig 1C), indicating that the hydrophobic environment drives the formation of smaller oligomers. His_6X_-Hsp20_KKK_ did not exhibit any change in hydrodynamic diameter with or without SDS and remained in a smaller form. To further analyze the protein size, we ran the purified recombinant proteins through a Superdex-200 column. The elution profile revealed that His_6X_-Hsp20 primarily forms higher-order oligomers, approximately 18-mer (Fig 1D). Consistent with previous findings, His_6X_-Hsp20 formed dimers upon treatment with 0.4 mM SDS (Fig 1D). His_6X_-Hsp20_KKK_, however, formed only dimers, with no higher-order oligomers (Fig 1D). We also examined the conformational changes associated with His_6X_-Hsp20 during its oligomer-dimer transition by measuring intrinsic tryptophan fluorescence. Interestingly, increasing the temperature led to a reduction in fluorescence (Fig 1E), suggesting that tryptophan residues become surface-exposed as the protein transitions from an oligomer to a dimer. Upon cooling, fluorescence intensity increased (Fig 1F), indicating a reversible transition. Similarly, SDS treatment reduced fluorescence in His_6X_-Hsp20 compared to the untreated protein (Fig 1G), confirming structural changes in response to temperature stress or a hydrophobic environment. These findings demonstrate the oligomeric plasticity of His_6X_-Hsp20. sHsps exhibit ATP-independent protein aggregation prevention activity. To assess whether *U. maydis* Hsp20 has similar activity, we performed a DTT-induced protein aggregation assay using lysozyme as the substrate. Aggregation was monitored by light scattering at 360 nm. Both His_6X_-Hsp20 and SDS-treated His_6X_-Hsp20 significantly reduced light scattering in the lysozyme-DTT mix, confirming their aggregation prevention activity (Fig 1H). His_6X_-Hsp20_KKK_ exhibited an activity comparable to His_6X_-Hsp20 (Fig 1H), suggesting that dimers are the active form of *U. maydis* Hsp20.

**Fig 1.**
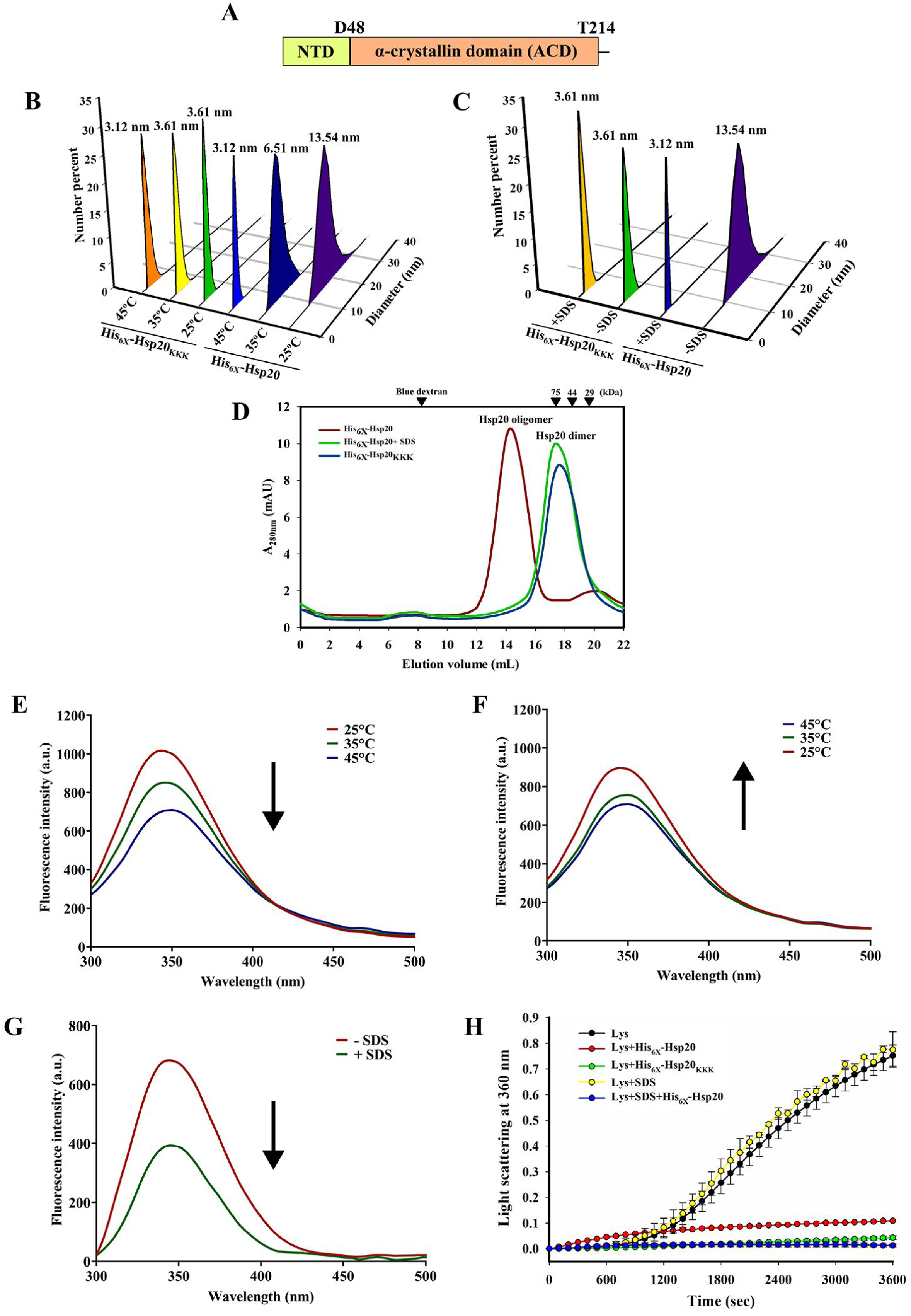
Oligomeric plasticity and protein aggregation prevention activity associated with *U. maydis* Hsp20. (A) Domain organization of Hsp20. (B & C) Graph showing Dynamic light scattering (DLS) data of His_6X_-Hsp20 and His_6X_-Hsp20_KKK_ at different temperature (B) and in the presence or absence of SDS (C). The diameter of the predominant oligomeric form is indicated at the top of each sample peak. (D) Gel filtration profile of His_6X_-Hsp20, SDS treated His_6X_-Hsp20 and His_6X_-Hsp20_KKK_ indicating the oligomer dimer equilibrium of the His_6X_-Hsp20 in the presence and absence of SDS. X axis represents elution volume fractions from the gel filtration column and Y axis represents absorbance at 280 nm. Black arrows mark the elution fractions representing indicated molecular weight markers. (E) & (F) Graph showing the change in intrinsic tryptophan fluorescence associated with His_6X_-Hsp20 with increasing (E) and decreasing (F) temperature. Arrows indicate the order of change in temperature, upward arrow indicates increasing and downward arrow indicates decreasing temperature. (G) Graph showing the change in the intrinsic fluorescence associated with His_6X_-Hsp20 in the presence and absence of SDS. Arrow indicates a decrease in the intrinsic fluorescence intensity in the presence of SDS. (H) Graph showing protein aggregation prevention activity associated with His_6X_-Hsp20 and His_6X_-Hsp20_KKK_ against DTT induced aggregation of lysozyme (Lys). Light scattering at 360 nm is represented by the Y axis and was used as the measure of protein aggregates formed. X axis represents the incubation time. Data represents average values calculated from at least three independent biological replicate experiments. Error bars represent standard deviation.

### Hsp20 deletion mutant of *U. maydis* exhibits aberrant colony and cellular morphology

In order to determine the cellular function of Hsp20 in *U. maydis*, we generated a deletion mutant of the respective gene in the haploid solopathogenic strain SG200. SG200Δ*hsp20* differed significantly from SG200 WT in both cellular and colony morphologies. While SG200 WT cells formed smooth and sticky colonies on PD agar media, the SG200Δ*hsp20* colonies appeared dry and exhibited reduced adherence to the agar surface (Fig 2 A). As perturbations in the colony morphology is a direct representation of changes in cellular morphologies, we compared the cells from the axenic broth culture of both the strains microscopically. Compared to the SG200 WT cells that show a characteristic cigar like morphology, SG200Δ*hsp20* cells exhibited more rounded appearance in the late exponential/stationary phase of growth (Fig 2B). A measurement of the relative cellular width of SG200Δ*hsp20* and SG200 WT cells revealed a significant increase in case of the deletion mutant (Fig 2C). In addition, compared to the SG200 WT cells, the sporidial cells of SG200Δ*hsp20* were found to remain attached to each other even after cell division resulting in the formation of long chains of cells. This indicates a possible defect in the cell separation events in case of SG200Δ*hsp20* (Fig 2D). Long chains of cells that fail to separate tend to entangle and aggregate in liquid cultures. Accordingly, the axenic culture of SG200Δ*hsp20* showed faster sedimentation compared to SG200 WT when allowed to stand still for 30 min (Fig 2E). Expression of a wild type copy of *hsp20* under the control of either a constitutive or the native promotor in the *hsp20* deletion background however reversed the phenotype completely. To find out whether the cell separation defect associated with SG200Δ*hsp20* is due to abnormal septation during cell division the cell division septa in the sporidial cells from both the wild type and mutant strains were visualised through calcofluor white staining. In case of SG200 WT two very closely spaced septa were noticed adjacent to one of the poles of the sporidial cells indicating proper budding events. On the contrary, the SG200Δ*hsp20* cells showed the presence of one or more septa positioned in the middle of the cells (Fig 2F). To further investigate any difference in the budding pattern of SG200Δ*hsp20* and SG200 WT cells we stained cells from both the strains with wheat germ agglutinin-Alexa Fluor 488 (WGA-AF 488). Like calcofluor white WGA also binds to cell wall components but exhibit a different staining pattern emphasising more on the regions of active deposition of cell wall components (Nagata & Burger, 1974). Since, budding involves active cell wall synthesis at the bud initiation sites, we compared the cell polarity during budding in SG200 WT and SG200Δ*hsp20* cells using WGA-AF 488 staining. Interestingly, the SG200Δ*hsp20* cells showed strong WGA-AF 488 staining at multiple sites within the cell walls including the two polar ends. This indicated a multipolar budding phenotype associated with the deletion mutant that is in sharp contrast to the wild type *U. maydis* cells that exhibit a unipolar budding (Fig 2G). We also tested for any abnormality in nuclear division in the mutant strain by staining the cells with 4’, 6-diamidino-2-phenylindole (DAPI). The SG200Δ*hsp20* cells however showed proper nuclear division and segregation between the mother and the daughter cells (Fig 2H). Septins are GTP-binding cytoskeletal components that are known to form hetero-oligomeric complexes to produce higher-order structures like collars, bands and filaments and are known to drive cytokinesis (Mostowy & Cossart, 2012). As SG200Δ*hsp20* cells showed defects in cytokinesis, we were curious to know whether the deletion of *hsp20* abrogates the sub-cellular localization of septin collars during cytokinesis. A previous study has shown that *U. maydis* codes for four septins (Sep1-4) and all these septins were seen to form collars at the bud neck to facilitate cell separation (Alvarez-Tabarés *et al.,* 2010). To visualize the septin collars we expressed a N-terminal eGFP tagged Sep2 in both SG200 WT and SG200Δ*hsp20* cells. As expected, in SG200 WT cells, GFP-Sep2 formed a single collar at the bud neck, indicating proper cytokinesis. Interestingly, although GFP-Sep2 could form collars in SG200Δ*hsp20* cells, their localization was altered. Instead of localising to the bud neck, we found septin collars present at multiple sites within the cell cortex of both the mother cell and the emerging bud (Fig 2I). As correct positioning of the septin collar is required for the separation of the bud from mother cell, the defects in cell separation in SG200Δ*hsp20* cells are probably due to the mislocalization of septin collars.

**Fig 2.**
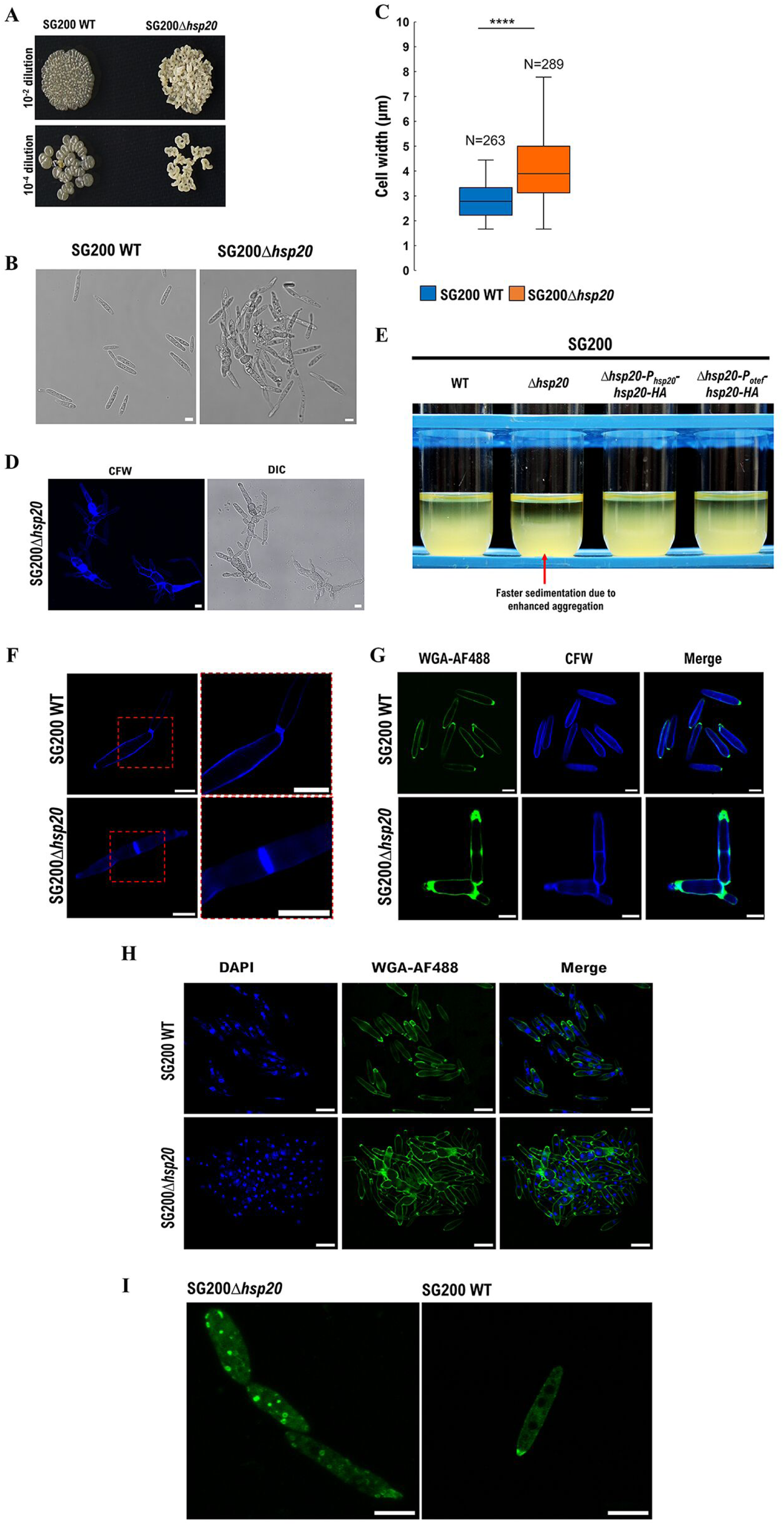
Growth properties and cellular morphology of *U. maydis* under axenic culture conditions. **(A)** Potato dextrose (PD) plates showing the colony morphology of SG200 WT and SG200Δ*hsp20*. The OD_600_ of exponentially growing cells were adjusted to 1.0 and spotted on PD plates in two different dilutions indicated. The plates were incubated at 28°C for 48 h before the image was taken. **(B)** Differential Interference Contrast (DIC) images showing cellular morphology of SG200 WT and SG200Δ*hsp20* captured in a confocal microscope. Scale bar corresponds to 5 µm. **(C)** Boxplot representing distribution of cell width in SG200 WT and SG200Δ*hsp20* cells. Widths of at least 80 cells from each of three independent experiments were measured and plotted in the graph. Statistical significance was determined using unpaired two-tailed T-test (****: P<0.0001). **(D)** Fluorescence micrographs of calcofluor white (CFW) stained cells of SG200Δ*hsp20*. Both fluorescence and DIC images are shown. Scale bars correspond to 5 µm. **(E)** Cell sedimentation assay of the indicated strains. Cultures of the indicated strains were adjusted to similar OD_600_ and were allowed to stand still for 30 min. SG200Δ*hsp20* cells sediment much faster due to enhanced aggregation of septate cells, as indicated by the appearance of zone of clearance at the top of the liquid cultures. **(F)** Fluorescence micrographs of SG200 WT and SG200Δ*hsp20* cells showing the partitioning cell wall septa formed during cell division (red box). Right panel shows zoomed in images pointing the absence of two closely positioned septa in *hsp20* deletion mutant. Scale bar corresponds to 5 µm. **(G)** Fluorescence micrographs of exponentially growing SG200 WT (upper panel) and SG200Δ*hsp20* (lower panel) cells stained with CFW and WGA-AF488. WGA stains regions of cell walls undergoing active growth. WGA staining is visible restricted to the growing tips in SG200 cells. Scale bars correspond to 5 µm. **(H)** Fluorescence micrographs of SG200 WT (upper panel) and SG200Δ*hsp20* (lower panel) cells stained with DAPI and WGA-AF488. *hsp20* deletion mutants are not defective in nuclear division as indicated by the presence of a single nucleus in each compartment of the septate cells. Scale bars correspond to 10 µm. **(I)** Fluorescence micrographs of SG200 WT and SG200Δ*hsp20* cells expressing eGFP-Sep2-HA. eGFP fluorescence in SG200 WT cells is seen to localize at one of the poles of the cells indicating the site of budding. On the contrary, in SG200Δ*hsp20 cells,* eGFP-Sep2-HA is seen assembled as collars at multiple sites on the cell cortex of both the mother and daughter cells. Scale bars correspond to 5 µm.

### Hsp20 influences organization of actin cytoskeleton in *U. maydis*

In *S. cerevisiae* polarised cell growth has been demonstrated to be dependent on the asymmetric distribution of actin cytoskeleton (Pruyne & Bretscher, 2000). Since deletion of *hsp20* led to loss of polarity of cell growth in *U. maydis* we became curious to find out whether Hsp20 affects the actin cytoskeleton in this fungus. We therefore visualised the actin cytoskeleton in both SG200 WT and SG200Δ*hsp20* cells by expressing an N-terminal eGFP tagged Lifeact probe within these cells. Lifeact is a 17 amino acid peptide that binds to peripheral actin patches as well as F actin cables in *U. maydis* (Riedl *et al*., 2008). In case of SG200 WT the Lifeact signal corresponding to actin patches were found to be concentrated only on one of the cell poles. On the contrary in SG200Δ*hsp20* equally populated actin patches were observed in both the cell poles (Fig 3A) indicating an active bidirectional growth. In addition, the number of fluorescent puncta observed in SG200Δ*hsp20* throughout the cells significantly outnumbered that observed in SG200 WT cells. This further indicated depolarization of actin in *U. maydis* in the absence of Hsp20. Significant differences were also observed in the positioning of the actin cables within the wild type and the mutant cells. Unlike SG200 WT where the actin cables appeared to extend from the mother to the daughter cell, in SG200Δ*hsp20* these cables were found to extend towards both the cell poles (Fig 3A, video S1 & S2). To further investigate the involvement of Hsp20 in modulating the actin assembly dynamics in *U. maydis* we tested the interaction between recombinant Hsp20 and actin using Fluorescence Resonance Energy Transfer (FRET) where actin was tagged with ALEXA 488 and His_6X_-Hsp20 with ALEXA-532. With increasing concentration of Hsp20, an increase in energy transfer was observed confirming the interaction, with a dissociation constant of 99.62±1.09 nM (Fig 3B, S2). The amino acid sequences of small heat shock proteins often possess regions of context dependent disorderness (Clouser *et al*., 2019). Many of the recent studies have shown that such proteins with intrinsically disordered regions have an increased propensity of forming liquid condensates and undergo liquid-liquid phase separation (LLPS) (Mukherjee & Schäfer, 2023, Borcherds *et al*., 2021). Therefore, to get deeper insights into the mechanism through which Hsp20 interacts with actin we evaluated any LLPS activity that might be associated with Hsp20. Analysis of the primary amino acid sequence of Hsp20 for regions of disorderness revealed stretches of intrinsically disordered regions (Fig S3). To validate this, we performed circular dichroism (CD) analysis on purified His_6X_-Hsp20, observing a significant random coil structure at room temperature (Fig 3C, D), supporting the presence of disorder. Upon heating or SDS treatment, the random coil content slightly decreased (Fig 3C-F). Given Hsp20’s ability to form higher-order oligomers, we hypothesized that it might undergo LLPS at higher concentrations. Indeed, purified His_6X_-Hsp20 exhibited increased light scattering at 360 nm in the presence of 10% and 15% PEG, indicating the formation of larger structures in presence of macromolecular crowding agent (Fig 3G). These aggregates were visualised as smooth spherical liquid droplets under confocal microscope (Fig 3H). Fluorescence recovery after photobleaching (FRAP) confirmed their liquid nature (Fig 3I). Based on these data, we generated a phase diagram of Hsp20 LLPS behaviour at varying protein and PEG concentrations (Fig 3J). We also assessed actin’s phase behaviour in the presence and absence of Hsp20. As expected, actin alone did not undergo phase separation, even at high PEG concentrations. However, in the presence of Hsp20, actin formed liquid condensates (Fig 3K), suggesting that Hsp20 LLPS droplets capture actin within their liquid phase through protein-protein interactions during phase separation.

**Fig 3.**
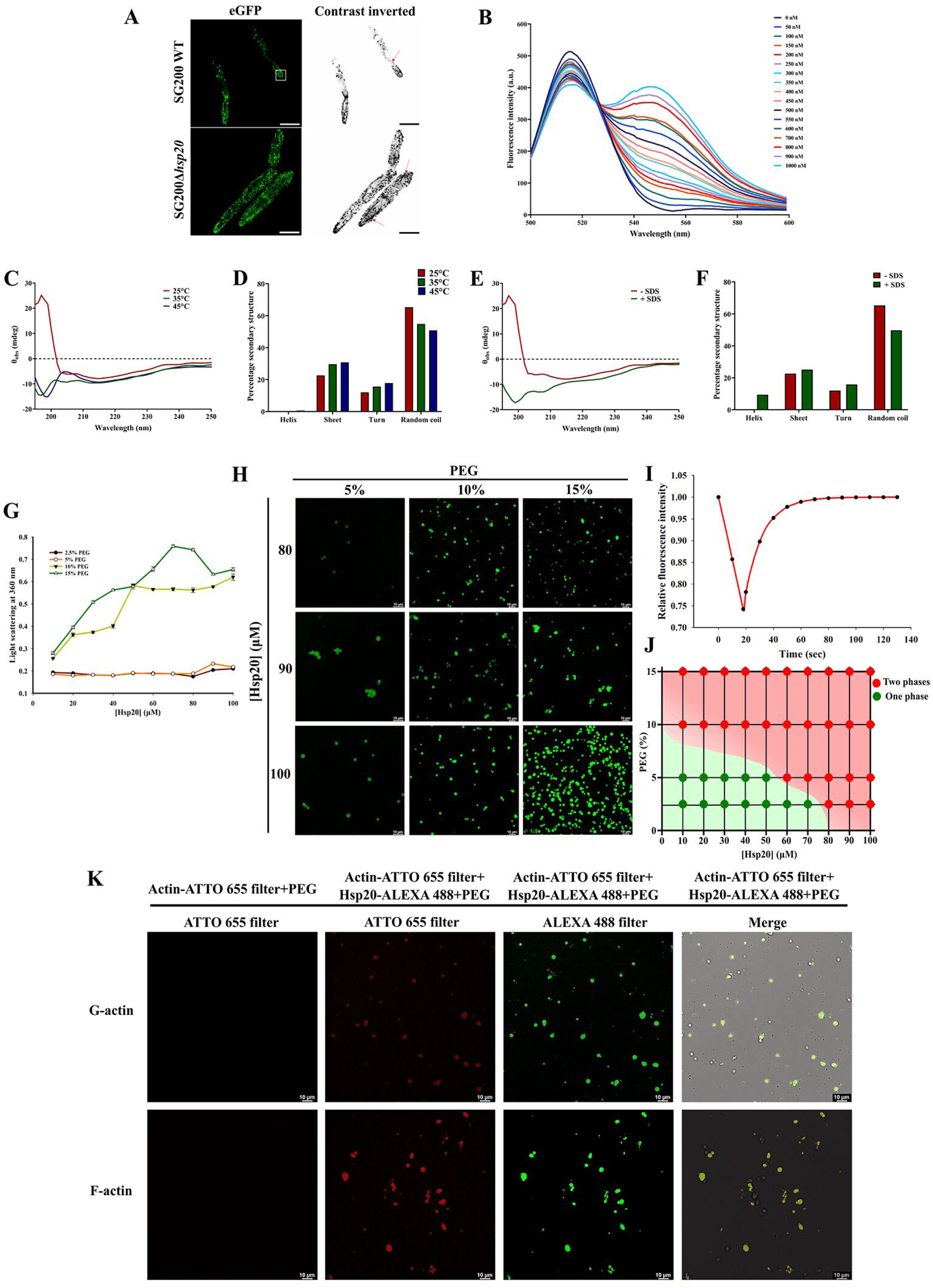
Liquid-liquid phase separation (LLPS) properties of Hsp20 and its ability to interact with actin. (A) Fluorescence micrographs showing the localization of actin in *U. maydis* SG200 WT and SG200Δ*hsp20* cells expressing eGFP-Lifeact-HA. Accumulation of actin patches at the tip of the growing cell is marked by white open square. Red arrows mark the actin cables. Right panels represent inverted-contrast images of the same cells as the left panels. Images shown are Z-projection of atleast 50 X-Y planes. Size bars correspond to 5 µm. (B) Positive Forster Resonance Energy Transfer (FRET) response showing physical interaction between Hsp20 tagged with Alexa 532 and actin tagged with Alexa 488. Samples were excited at 490 nm and emissions were recorded between 500 nm to 600 nm. (C-F) Circular dichroism (CD) spectral properties of Hsp20 at different temperatures (C, D) and in the presence of SDS (E, F) indicating significant proportion of random coil present that decreases slightly with external environmental factors. (G) Graph showing induced aggregation of Hsp20 with increasing concentration of the protein and polyethylene glycol (PEG). Protein aggregation was measured as light scattering at 360 nm. (H) Confocal microscopy images of ALEXA 488 tagged recombinant Hsp20 showing smooth spherical droplets when incubated in the presence of indicated concentration of PEG at 37°C for 1 h. (I) Graph showing fluorescence recovery after photobleach (FRAP) on the Alexa 488 tagged Hsp20 spherical droplets observed under a confocal microscope. (J) Phase diagram showing the existence of Hsp20 in two phases at concentrations above 60 µM Hsp20 and 5% PEG. (K) Fluorescence micrographs showing Hsp20 LLPS droplets that also absorb actin within the liquid phase through protein-protein interaction while undergoing phase separation. Hsp20 and actin were tagged with ATTO 655 and Alexa 488 tags respectively. Upper panels show interaction with G actin and lower panels show interaction with F actin.

### Hsp20 partially colocalises with actin and septin in *U. maydis* sporidial cells

To determine the contribution of Hsp20 in the actin cytoskeletal dynamics in *U. maydis* sporidial cells, we visualised Hsp20 in the context of actin distribution in SG200-[*mcherry-hsp20-HA*] [*eGFP-Lifeact-HA*]. The SG200-[*mcherry-Hsp20-HA*] [*eGFP-Lifeact-HA*] strain expressed an N-terminal mcherry and C-terminal HA tagged version of Hsp20 from *hsp20* genomic locus in the SG200-*eGFP-Lifeact-HA* background. When observed under a confocal microscope the cells showed partial colocalization of Hsp20 with actin patches but not with the actin cables (Fig 4A). A very similar pattern of fluorescence signal colocalization was also noticed in case of eGFP-Sep2-HA and mcherry-Hsp20-HA in SG200-[*mcherry-Hsp20-HA*] [*eGFP-Sep2-HA*]. Like in case of actin, Hsp20 did not colocalise with polymeric structures of septin assembled on the cell cortex. In non-dividing cells colocalization of Hsp20 and Sep2 was observed in cytosolic punctate structures (Fig 4B). Interestingly in a dividing cell, where Sep2 localization was observed both in cytosol and at the bud neck in the form of collar, mcherry signal from mcherry-Hsp20-HA was found only at the cytosolic Sep2 puncta (Fig 4C).

**Fig 4.**
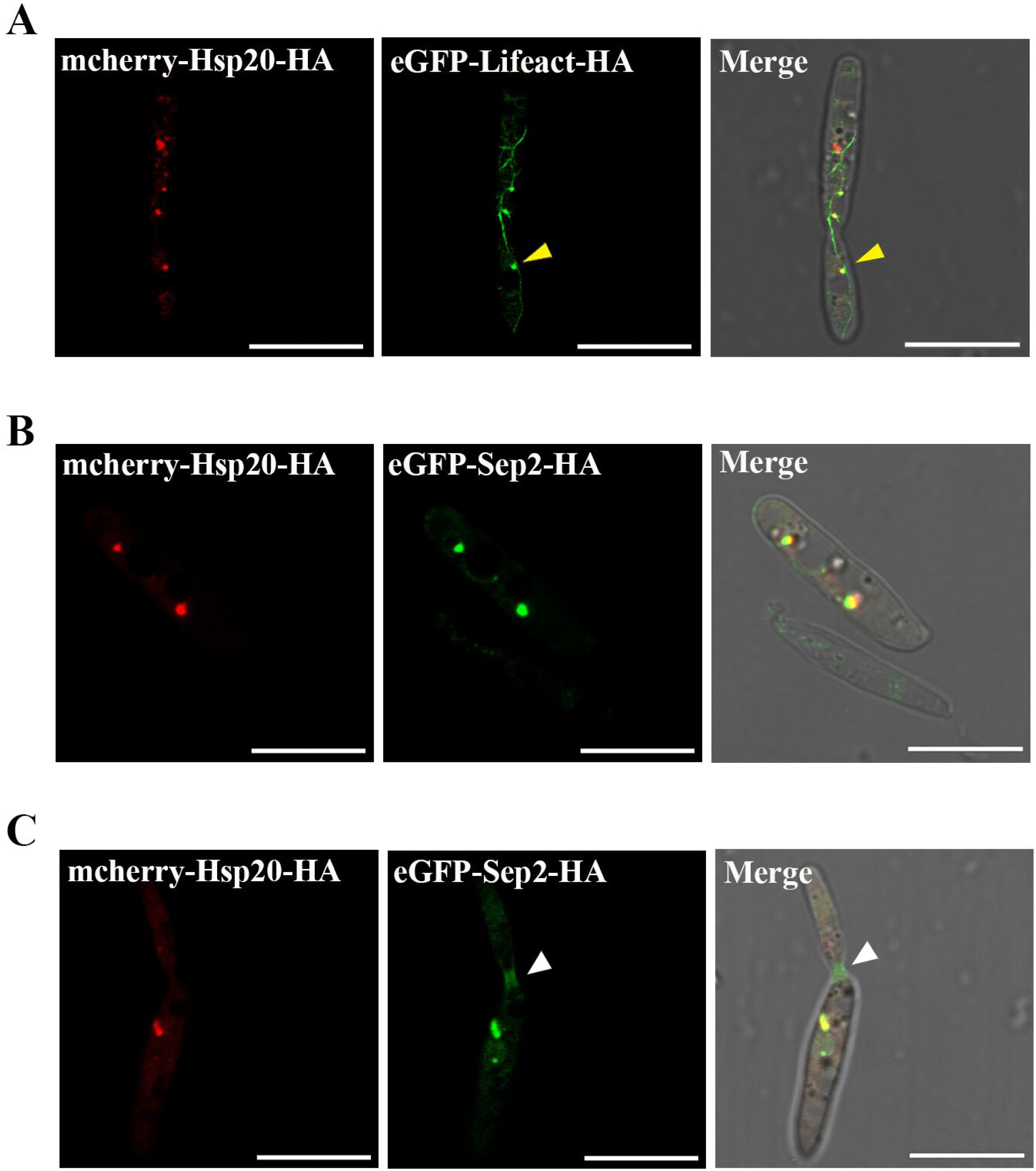
Cellular localization of Hsp20 in the context of actin and septin. Confocal microscopy images of the sporidial cells of *U. maydis* SG200-[*mcherry-hsp20-HA*] [*eGFP-Lifeact-HA*] (A) and SG200-[*mcherry-hsp20-HA*] [*eGFP-Sep2-HA*] (B & C) strains co-expressing mcherry-Hsp20-HA/eGFP-Lifeact-HA and mcherry-Hsp20-HA/eGFP-Sep2-HA respectively. Yellow arrow heads mark actin patches. White arrow heads show Sep2 collar in the bud initiation site. Size bars indicate 10 µm.

### Hsp20 influences endocytosis in *U. maydis*

Previous studies have established a connection between the organization and movement of actin patches with the dynamics of the endocytosis process (Mooren *et al*., 2012). Since the organization of the actin cytoskeleton is significantly influenced by Hsp20 in *U. maydis* we further investigated whether *SG200Δhsp20* can execute proper endocytosis by monitoring the uptake of an amphiphilic styryl dye FM4-64 in a pulse-chase experiment. When compared to similarly treated SG200 WT, the mutant cells resembled the wild type cells in exhibiting negligible retention of the fluorescent signal in the plasma membrane after a chase period of 45 mins. In both the cases the fluorescent signal was observed primarily within the vacuolar membranes (Fig 5A). However, unlike SG200 WT that exhibited about 4-5 vacuoles per cell, SG200Δ*hsp20* showed numerous vacuoles with relatively smaller diameter than that observed in the wild type cells (Fig 5A). A quantification of the relative number and size of the vacuoles in SG200 WT and SG200Δ*hsp20* showed about 4-5 times more vacuoles in the deletion mutant that are about half the size of those observed in the wild type cells (Fig 5 B& C). To further confirm we also stained the vacuoles in both SG200 WT and SG200Δ*hsp20* with CellTracker^TM^Blue CMAC (7-amino-4-Chloromethylcoumarine) dye that localises within the vacuolar lumen. CMAC staining also showed increased number of endocytic vacuoles and with reduced diameter in SG200Δ*hsp20* cells when compared to the wild type cells (Fig 5D).

**Fig 5.**
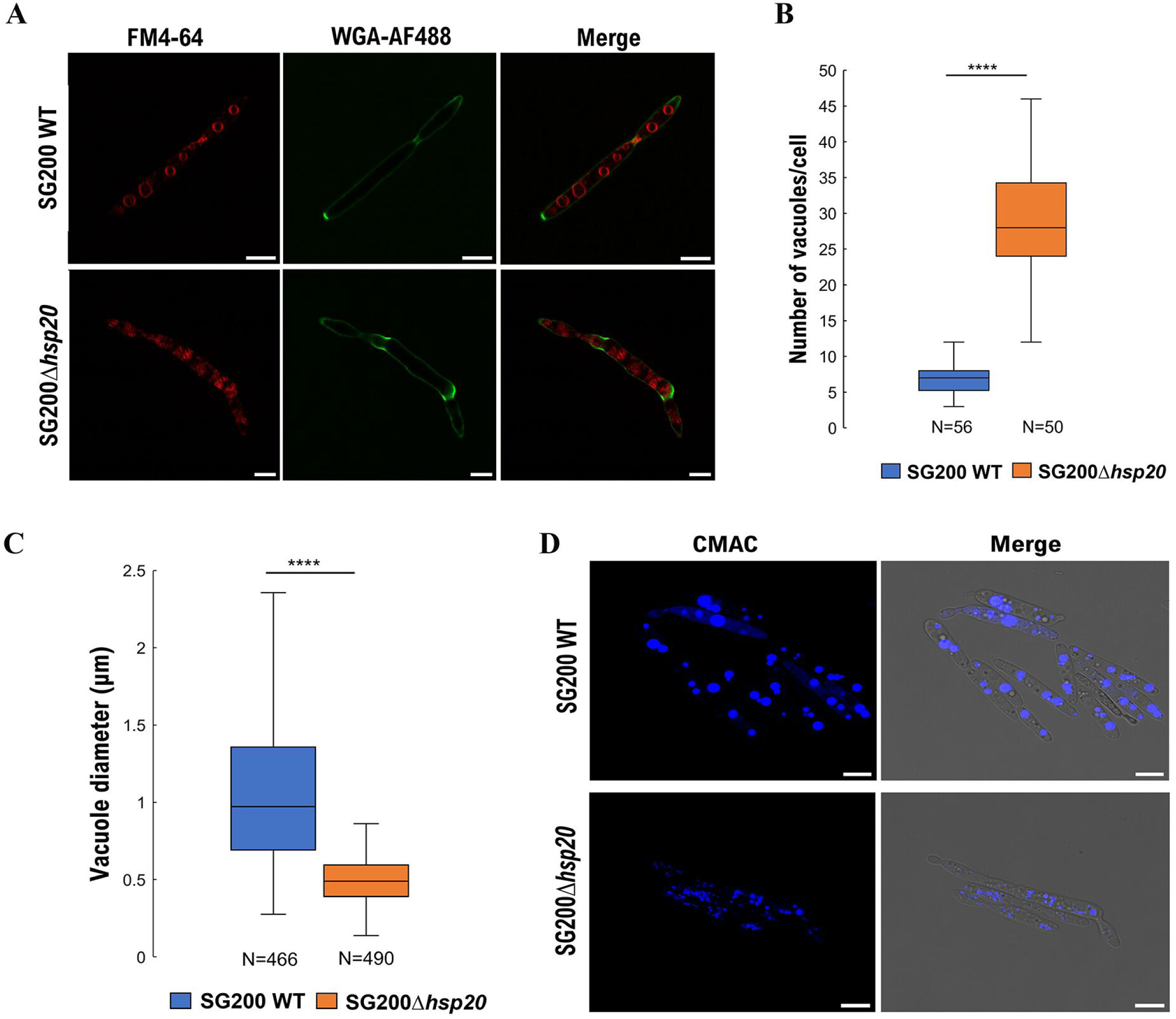
Endocytosis in *U. maydis* SG200 WT and SG200Δ*hsp20*. **(A**) Fluorescence micrographs of exponentially growing SG200 WT and SG200Δ*hsp20* cells that were pulsed with FM4-64 for 5 min and chased for 45 min. Cell walls were stained with WGA-AF488. A significantly increased number of smaller vesicles are observed in SG200Δ*hsp20* cells compared to the wild type cells. Scale bars correspond to 5 µm. (B & C) Boxplot representing vacuolar number per cell **(B)** and vacuolar diameter **(C)** of indicated strains. SG200Δ*hsp20* cells exhibit higher vacuolar number with reduced diameter when compared to SG200 WT cells. Vacuolar number and diameter were measured from three independent experiment. Statistical significance was calculated using unpaired two-tailed T-test (****: P<0.0001). **(D)** Fluorescence micrographs showing SG200 WT and SG200Δ*hsp20* cells stained with 7-amino-4-chloromethylcoumarin (CMAC) dye that stains vacuolar lumen. Scale bars correspond to 5 µm.

### Hsp20 is involved in the mating and filamentation processes of *U. maydis*

As deletion of *hsp20* dramatically influences cellular morphology in *U. maydis*, we were curious to find out whether the deletion mutant can undergo the dimorphic switch from apathogenic sporidial form to the pathogenic filamentous form. We therefore spotted the SG200Δ*hsp20* and SG200 WT on PD agar plates supplemented with charcoal that mimics the maize leaf surface and induces filamentation. *U. maydis* SG200 is a solo-pathogenic strain that harbours genes coding for a functional bE1/bW2 heterodimer and constitutively expresses the pheromone of opposite mating type (mfa2). Thus, SG200 despite being a haploid strain can undergo filamentation in the absence of a mating partner and can be used for studying sporidia to filament transition in *U. maydis* (Kämper *et al*., 2006, Bölker *et al*., 1995). Induction of filamentation on charcoal plates is observed as white fuzzy growth due to the formation of aerial hyphae as opposed to the smooth and sticky colonies representing the sporidial cells. Interestingly, SG200Δ*hsp20* showed considerably reduced filamentation compared to SG200 WT when grown on PD agar media supplemented with charcoal (Fig 6A). The reduced filamentation phenotype associated with the deletion mutant was also found to be well complemented by a wild type copy of the *hsp20* gene expressed either under the control of the native or a constitutive promotor (otef) from the *ip* locus of SG200Δ*hsp20* (Fig 6A). When observed under a microscope majority of the SG200Δ*hsp20* cells from PD charcoal plates were found to be arrested in the sporidial stage. Nevertheless, few filaments were also noticed among the mutant population, but the length of the filaments were found to be significantly shorter than that observed in case of the SG200 WT cells (Fig 6B). To further assess the involvement of Hsp20 in regulating the dimorphic switch in *U. maydis*, we compared the expression of *hsp20* between the sporidial and filamentous cells of SG200 WT strain grown on PD agar plates supplemented without or with charcoal respectively. As expected, the expression of *hsp20* was found to be induced when the cells were grown on PD charcoal plates (Fig 6C). This indicated that Hsp20 is indeed required for the dimorphic switch in the fungus. In *U. maydis* the morphological transition from sporidia to filament is linked to mating (Banuett & Herskowitz, 1994) (Kahmann *et al*., 1995). Therefore, we further tested any involvement of Hsp20 in the mating process of *U. maydis* using *hsp20* deletion mutants generated in the mating compatible strains FB1 and FB2. Accordingly axenic cultures of wild type and *hsp20* deletion mutant strains of *U. maydis* of opposite mating types were mixed in equal proportions and spotted on solid PD charcoal media. The inoculated agar plates were incubated for the appearance of white fuzzy growth representing filamentation due to mating. Interestingly a significantly reduced mating was observed in case of FB1Δ*hsp20* X FB2Δ*hsp20* and FB1Δ*hsp20* X FB2 WT but not in case of FB1 WT X FB2Δ*hsp20* (Fig 6D). Mating and subsequent filamentation in *U. maydis* is initiated by the recognition and binding of a cell surface pheromone receptor on one of the mating partners with the pheromone released from the other partner. This interaction between the pheromone and its receptor leads to the phosphorylation and activation of a major transcription factor Prf1 via two different signalling cascades, the MAPK and the cAMP. The Prf1 in turn induces the expression of both the ‘a’ type genes coding for pheromone/pheromone receptor and the ‘b’ type genes coding for the bE/bW transcription factor, the master regulator of pathogenic gene expression and filamentous growth in *U. maydis* (Brachmann et al., 2001). To test whether the reduced filamentation upon *hsp20* deletion in *U. maydis* was due to decreased efficiency of mating signal transduction the relative expression of *prf1*, *bE1* and *bW2* were compared between SG200 WT and SG200Δ*hsp20* following 24 h growth on PD charcoal plate. All the three genes showed reduced expression in the deletion mutant compared to the wild type strain indicating possible involvement of Hsp20 in the downstream signal transduction following pheromone-receptor interaction (Fig 6E). The reduction in the observed filamentation associated with SG200Δ*hsp20* therefore can be attributed to the decreased bE/bW heterodimer levels. To further support this observation, we studied sporidia to filament transition of wild type and *hsp20* deletion mutant in *U. maydis* AB33 strain. AB33 cells possess *bE and bW* genes under the control of a nitrate inducible promotor. Thus, when AB33 cells are grown in a nitrate minimal media the *b* genes are induced independent of the pheromone/pheromone-receptor signalling pathway and lead to morphological transition of the cells from sporidia to filaments. Interestingly AB33Δ*hsp20* did not show any defect in sporidia to filament transition when transferred to a nitrate minimal media. The percentage of filament formation in case of AB33Δ*hsp20* was comparable to that in case of the wild type AB33 strain (Fig S4). As SG200Δ*hsp20* showed defective filamentation we were curious to find out whether the fungus is capable of completing its life cycle marked by the formation of viable spores in the absence of Hsp20. Accordingly, *Z. mays* plants infected with either SG200 WT or SG200Δ*hsp20* were monitored for the progression of infection. The infected leaves were stained with Wheat Germ Agglutinin-Alexa Fluor 488 (WGA-AF488) and propidium iodide (PI) that stain fungal and plant cell walls respectively (Redkar *et al*., 2018). While majority of the SG200 WT cells formed long penetrative hyphae by 2 dpi, very few short hyphae were observed in case of SG200Δ*hsp20*. Moreover, the deletion mutant also exhibited substantially reduced hyphal proliferation compared to the wild type cells at 4 dpi. In addition, the hyphae formed by SG200Δ*hsp20* did not show any fragmentation or aggregation within gelatinous matrix that finally leads to the formation of mature teliospores at around 12 dpi (Fig 6F). The inability of SG200Δ*hsp20* to sporulate and complete pathogenic lifecycle is also accompanied by its attenuated virulence as assessed by the relatively lesser number of tumors formed on the leaf surfaces (Fig 6G). Moreover, the ligula of SG200Δ*hsp20* infected plants did not show the characteristic swelling upon infection and were also found to be devoid of spore masses (Fig 6H). To further confirm the lack of sporulation in SG200Δ*hsp20*, we compared the expression of *ros1* between *hsp20* deletion mutant and the wild type strain of *U. maydis* SG200 at different stages of infection. Ros1 is the major regulator of sporogenesis and expression of ‘late effectors’ in *U. maydis*. Previous studies have shown that induced expression of *ros1* during the later stages of infection is essential for karyogamy, hyphal aggregate formation and sporogenesis in *U. maydis* (Tollot et al., 2016). While SG200 WT cells showed increased expression of *ros1* in the later stages of infection with the highest expression at 12 dpi, SG200Δ*hsp20* did not exhibit any induction in expression even at 12 dpi (Fig 6I). Thus, the lack of Hsp20 in *U. maydis* not only reduces the efficiency of the fungus to form hyphae but also interferes with the progression of infection and attenuates virulence.

**Fig 6.**
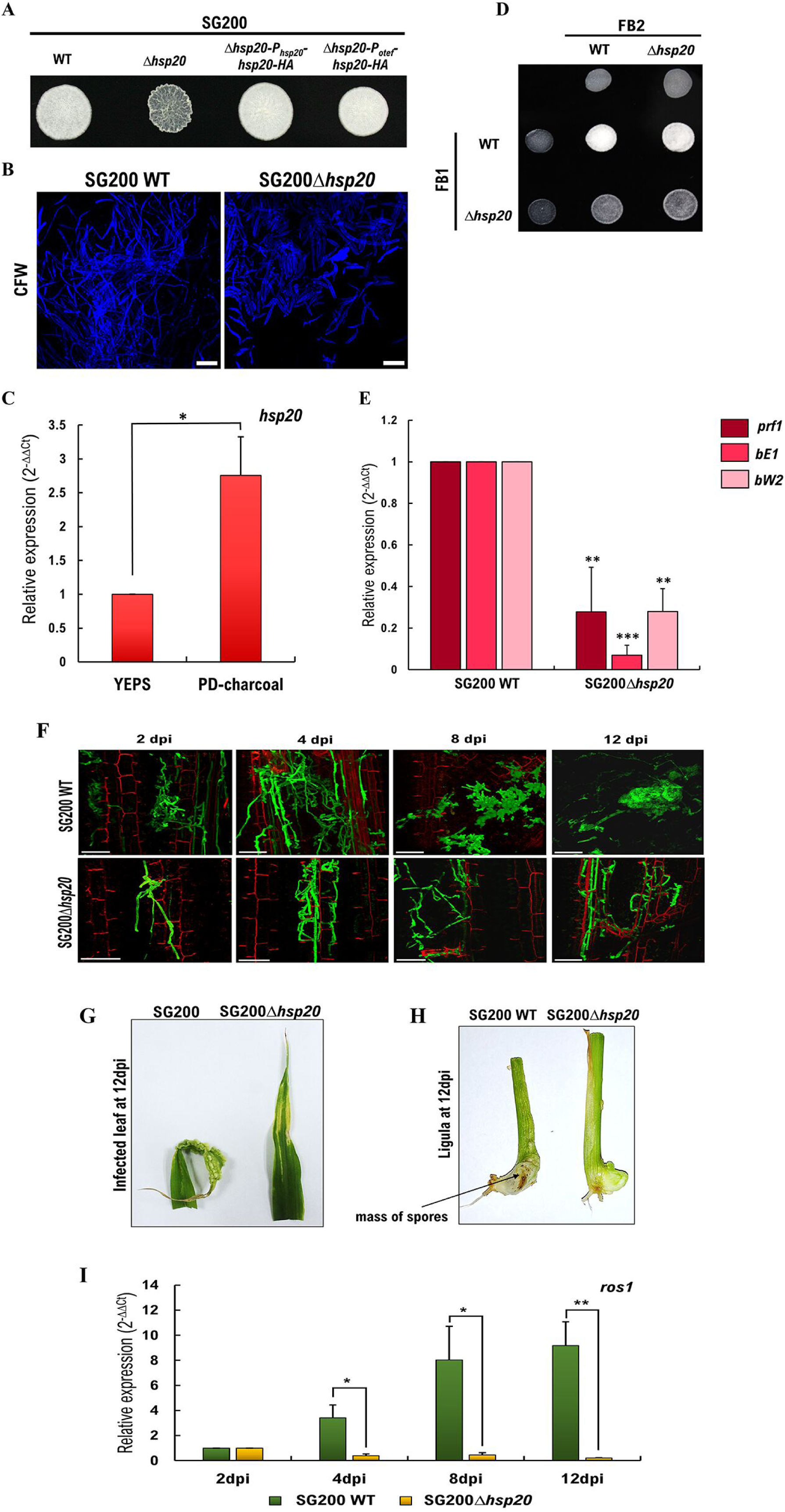
Filamentation and pathogenicity in *U. maydis* SG200 WT and SG200Δ*hsp20*. **(A)** PD-charcoal agar plates showing colony morphology of the indicated strains of *U. maydis.* Lack of fuzzy growth indicates reduced filamentation in *hsp20* deletion mutant compared to the wild type and the complementation strains. **(B)** Fluorescence micrographs of CFW stained SG200 WT and SG200Δ*hsp20* cells growing on PD-charcoal plates for 24 h. Scale bar corresponds to 10 µm. **(C)** Graph showing the relative expression of *hsp20* measured using qRT-PCR in SG200 WT cells grown in PD charcoal and YEPS media. Cells from PD charcoal plates were harvested by scraping off fungal growth from the agar surface after 24 h incubation at 28°C. The expression value was quantified with respect to exponentially growing SG200 WT cells in liquid YEPS medium. **(D)** Mating assay between sexually compatible *U. maydis* haploid cells. Cells from each strain were either spotted alone or in combination (as indicated) on PD charcoal plates and incubated for 24 h at 28°C. Appearance of white fuzzy growth is indicative of successful mating and consequent formation of filaments. **(E)** Graph showing the qRT-PCR analysis of transcription factors *prf1*, *bE1*, and *bW2* expression in SG200Δ*hsp20* cells, 24 h post-induction of filamentation. The relative expression values were calculated with respect to SG200 WT cells, 24 h post-induction of filamentation. Exponentially growing sporidial cells of indicated strains were spread on PD-charcoal plates and incubated at 28°C for 24 h. Cells were harvested by scraping off the growth from the agar surface and RNA was isolated. **(F)** Confocal microscopy images showing maize leaves infected with indicated strains of *U. maydis*. Leaf tissues were stained with WGA-AF488 (green, fungal cell walls) and propidium iodide (red, plant cell walls) to visualise different stages of development of the fungus. At least four-leaf samples from two independent experiments were analysed for each infection timepoint. Images shown are Z-projections of at least 100 X-Y planes. Scale bars correspond to 50 µm. **(G)** Leaves from SG200Δ*hsp20* infected seedlings showing reduced formation of large tumours at 12 dpi compared to the wild type infected ones. **(H)** Ligules of maize plants infected with indicated strains of *U. maydis* showing swelling and black spore mass in SG200 WT infected ligules but not in SG200Δ*hsp20*. **(I)** Graph showing the qRT-pCR analysis of *ros1* expression in leaf samples infected with either SG200 WT or SG200Δ*hsp20* strains, 4, 8, and 12 dpi. The expression value in SG200 WT & SG200Δ*hsp20* was calculated with respect to their respective expression at 2 dpi. Expression data in (C), (E) and (I) were normalized with respect to the expression levels of constitutively expressed gene, *ppi* (UMAG_03726). The data shown in all of these graphs present mean values calculated from three biological replicates. Error bars indicate standard error of mean. Statistical significance was calculated using unpaired two-tailed T-test (*: P<0.05, **: P<0.01, ***:P<0.001).

## Discussion

sHsps are important stress proteins that contribute significantly in cellular survival under conditions of proteotoxic stress. Misfolded proteins within a cell are sequestered in near native conformation by sHsps (Ungelenk *et al*., 2016). The molecular mechanism underlying the function of sHsps involve their ability to oligomerize. Some of the sHsps have also been found to phase separate within a cell owing to their ability to form higher order oligomers. Hsp40 protein, Hdj1 for instance was demonstrated to form LLPS both independently and together with an RNA binding protein FUS. This study showed that by forming co-LLPS with FUS Hsp40 keeps the protein in a condensed phase-separated state and inhibits it to form toxic amyloid fibril (Gu *et al*., 2020). In the present study we have shown LLPS property associated with *U. maydis* Hsp20. Furthermore, we also found induced LLPS formation in G-and F-actin in the presence of Hsp20 in in-vitro experiments. Interaction of actin with sHsps have been demonstrated earlier as well in few instances. For example in H9C2, rat cardiomyoblast cell line αβ crystalline was found to interact with actin filaments and protect its disorganization when treated with actin depolymerizing agent, cytochalasin B under conditions of heat stress (Singh *et al*., 2007). Likewise Hsp27/HspB1 monomers have been shown to bind F-actin filament along the side (Graceffa, 2011) as well as at the barbed ends (Benndorf *et al*., 1994). In a separate study non phosphorylated Hsp27 was found to bind and sequester purified actin monomers thereby making them unavailable to form filaments (During *et al*., 2007). Thus, together these studies indicate an important role of sHsps in the actin filament dynamics within a cell. In alignment with these findings *U. maydis hsp20* deletion mutant exhibited an altered layout of actin filaments and patches when compared to the wild type cells. In the wild type cells, one of the poles that serves as the origin of the daughter cell showed maximum density of the actin patches. On the contrary both the poles of the deletion mutant were equally populated with actin patches indicating a loss of polarity during budding. When the localization of Hsp20 within a *U. maydis* cell was traced the protein was found in distinct puncta (Fig S5). Furthermore, some of these puncta showed colocalization with eGFP signal in a *U. maydis* cell co-expressing mcherry-Hsp20-HA and eGFP-Lifeact-HA. Interestingly, we never obtained mcherry-Hsp20-HA signal in the actin cables in these cells. It is possible that Hsp20 sequesters a population of actin within a *U. maydis* cell and regulates its polymerization dynamics with the help of other factors that are yet to be revealed. In addition to actin, *hsp20* deletion mutant also exhibited aberrant positioning of septin collars within the cellular cortex. These proteins play important role in cell division and determination of cell polarity. In *S. cerevisiae*, septins form ring structures and localize within the cortex at the site of bud emergence. As the bud expands the ring structure transforms into an hour-glass structure and finally the daughter cell splits away from the mother cell (Marquardt *et al*., 2021). In the present study we demonstrate that in the absence of Hsp20, *U. maydis* septins assemble as ring structures throughout the cortex of the cells. This is in sharp contrast to the wild type cells where the rings localize specifically to one of the poles. Interestingly we found no association of Hsp20 with the septin polymers on the cell cortex through colocalization studies. However, like actin we did find colocalization of eGFP and mcherry signal in distinct puncta within the cytoplasm of *U. maydis* cell expressing mcherry-Hsp20-HA and eGFP-Sep2-HA simultaneously. We believe that like actin, Sep2 is also one of the proteins that is sequestered within Hsp20 punctate structures. It is likely that Hsp20 is involved in the molecular mechanism determining the recruitment and positioning of polymeric septin structures into the cortex of a cell at the site of bud initiation. Although not much is yet known about septin recruitment and positioning within a budding cell cortex there are few studies indicating important role of proteins like a small rho GTPase, Cdc42 and its effector Gic2 (Sadian *et al*., 2013). Any involvement of Hsp20 in modulating these proteins thereby influencing the correct positioning of septin structures at the cell cortex in *U. maydis* is elusive and awaits further investigation. Absence of Hsp20 protein in *U. maydis* was also found to have significant influence on the cellular endocytosis process. The increased number of endocytic vesicles with reduced diameter observed in SG200Δ*hsp20* compared to the wild type cells might be attributed to the membrane interaction properties of sHsps. One of the possibilities to describe the vesicle phenotype associated with the *hsp20* deletion mutant is that Hsp20 can facilitate vesicular fusion. Although, till date there are no such studies available that explored sHsps in the context of vesicular fusion. Nevertheless, in one study a positive influence of Hsp70 on the clathrin mediated endocytosis process in human hepatoblastoma cells was shown. However, the underlying mechanism involved in this process was proposed to be an accelerated internalization of receptor ligand complex and increased recycling of the receptor (Vega *et al*., 2010). Endocytosis is one of the cellular processes that is critical for the pathogenic development of *U. maydis*. Following mating between the compatible sporidia of the fungus, the pheromone/pheromone receptor complex is internalized through endocytosis initiating a cascade of signalling events leading to the activation of the bE/bW transcription factor (Fuchs *et al*., 2006). A defective endocytosis therefore is most likely to affect the mating process and subsequent downstream signalling in *U. maydis.* Consistent with this hypothesis we found a defective mating between FB1Δ*hsp20* and FB2Δ*hsp20*. Furthermore, we also observed significant reduction in the induction of filamentation in SG200Δ*hsp20* cells when compared to the wild type cells when grown on charcoal agar media. This defect became even more clear when the *hsp20* deletion mutant failed to complete its pathogenic lifecycle within the infected *Z. mays* with absolutely no spores formed. This study therefore presents Hsp20 as an important regulator of multiple cellular pathways summarised in Fig 7 that lead to the pathogenic development of *U. maydis*. However, further study is needed to determine the mechanistic differences in the activity of Hsp20 in the regulation of these pathways.

**Fig 7.**
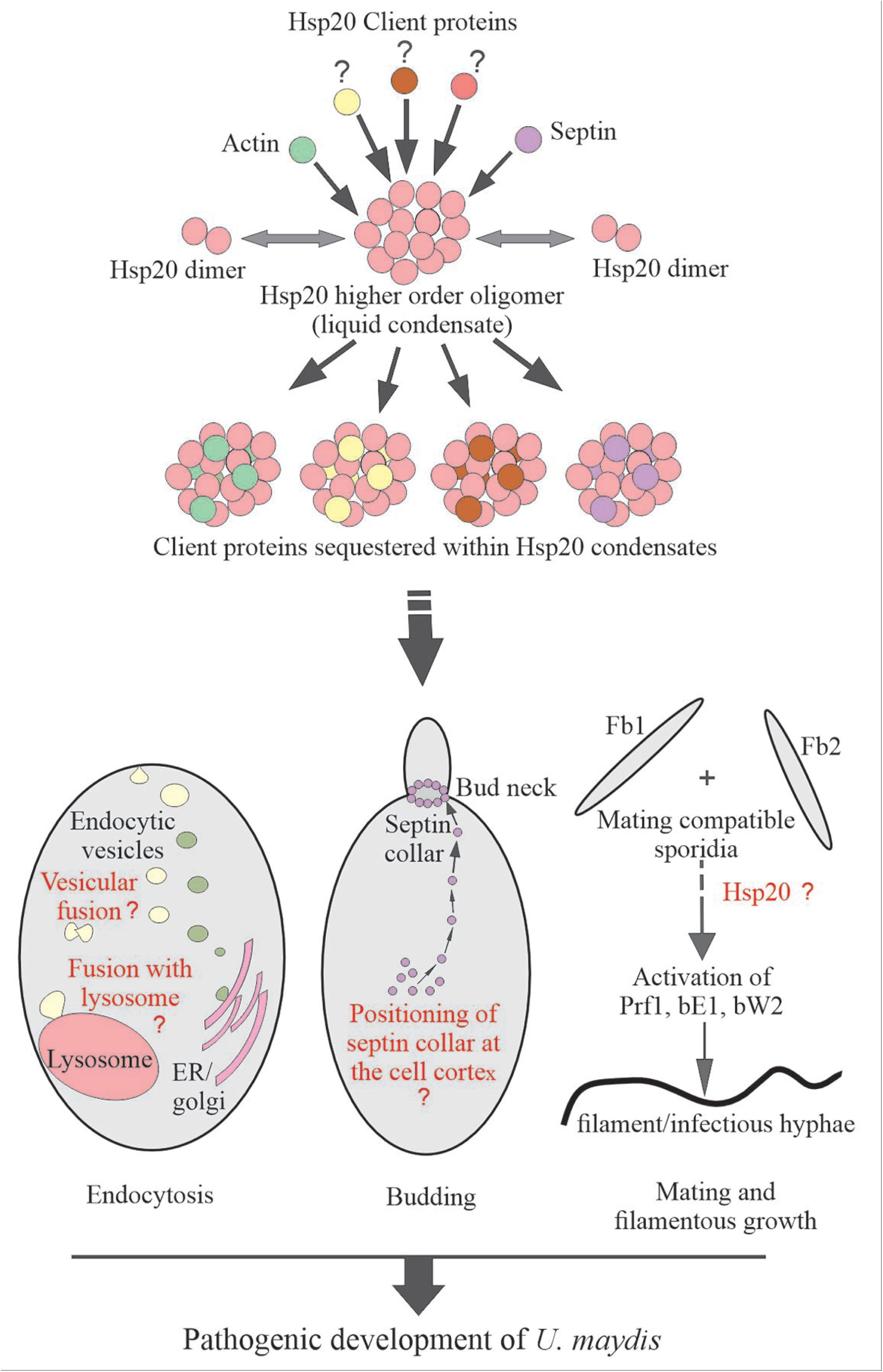
Model depicting possible ways of Hsp20 functioning in *U. maydis*. Under in vitro condition Hsp20 remains in dynamic equilibrium between dimer and higher order oligomers. The higher order oligomers exhibit phase separation in the presence of a crowding agent and form liquid condensates. In the presence of Hsp20 client proteins, Hsp20 can form co-LLPS leading to inclusion of the client proteins within the condensates. We have shown such co-LLPS formation of Hsp20 with actin in this study. Besides actin other client proteins might include septins like Sep2 that we have found to interact with actin within *U. maydis* cells. Within a cell, inclusion of client proteins into Hsp20 condensates leads to their sequestration and thereby limited or regulatable availability for cellular function like budding and mating in *U. maydis*. Hsp20 is involved in the proper positioning of septin collar at the bud neck and thereby determining polarity during cell division. During mating, Hsp20 is involved in the regulation of mating signal and subsequent filamentation. Hsp20 might also be involved in vesicular fusion among the endocytic vesicles and between endocytic vesicles and lysosomes during endocytosis. All these cellular functions together direct pathogenic development of *U. maydis*.

## Methods

### Generation of U. maydis strains

All the *U. maydis* strains were generated using protoplast transformation with linearised vectors. For the protoplastation of different background strains the respective strains were grown in YEPSL media (0.4% sucrose, 0.4% peptone, 1% yeast extract) till OD_600_ reached 0.8. At this stage the cells were washed once with SCS buffer (20 mM Tris sodium citrate dihydrate, 1 M D-sorbitol, pH 5.8) and then incubated in SCS buffer (20 mM trisodium citrate dihydrate, 1 M D-sorbitol, pH 5.8) supplemented with 12.5 mg/ml Lysing enzyme (Sigma) on ice until atleast 60% protoplastation was achieved. The cell wall lysis was finally stopped by adding 5 reaction volumes of SCS buffer. The resulting cell suspension was then centrifuged at 500 g for 10 min at 4℃. The cell pellet thus obtained was washed once with STC buffer (10 mM Tris HCl, pH 7.5, 100 mM CaCl_2_ and 1 M D-sorbitol) and finally stored in fresh STC buffer in 50 μl aliquots at -80℃ till further use. In order to carry out transformation of the prepared protoplasts 1 μl of 15 mg/ml heparin solution was added to 20 μl of the linearized p123 derived plasmids or the *hsp20* knock out cassette. To this DNA preparation 50 μl of respective *U. maydis* protoplasts were added and the final suspension was incubated on ice for 10 min. Following this 500 μl of 50 % solution of PEG 4000 in STC buffer was added to the protoplast and the resulting suspension was incubated on ice for another 15 min. After this the protoplast suspension was spread on regeneration agar (1 % Yeast extract, 2% peptone, 2 % sucrose and 1M sorbitol in water) supplemented with appropriate antibiotics and incubated at 28℃ till visible colonies were formed. Positive transformants were screened using polymerase chain reaction (PCR) based method and confirmed by Southern blot analysis. All the *U. maydis* strains used in this study are enlisted in Table S1. The cloning strategies for generating different plasmid constructs are enlisted in Table S2.

### PCR based screening of U. maydis transformants

In order to screen for the *U. maydis* transformants with insert integration within the succinate dehydrogenase (cbx) locus, PCR was performed on the genomic DNA from the transformants with the insert specific forward primer and the p123 vector backbone specific reverse primers (3_p123). For the transformants with N terminal genome tagging, PCR was carried out with tag specific forward primer and a reverse primer from the right border of the gene. For the screening of *hsp20* deletion mutant three sets of PCR were performed for each candidate transformant with primers specific to the hygromycin resistance cassette (5_hygR/3_hygR), the gene locus (5_hsp20_LB/3_hsp20_RB) and the gene ORF (5_hsp20/3_hsp20). All the primers used in this study are enlisted in Table S3.

### Genomic DNA isolation from U. maydis

Genomic DNA isolation from *U. maydis* was carried out as described previously (Sambrook & Russell, 2006) with some modifications. The sporidial cells from the overnight axenic culture of different *U. maydis* strains were lysed in 1 ml of a 1:1 (v/v) solution of gDNA isolation buffer (150 mM NaCl, 200 mM Tris-HCl pH 8.0, 10 mM EDTA, 1% SDS) and phenol, chloroform, isoamylalcohol mixture in the ratio 25:24:1 (v/v) (Himedia) using bead beating method with 450 μm – 650 μm acid washed glass beads (Sigma). Following lysis, the suspension was centrifuged at 16000 g for 15 min at room temperature to facilitate phase separation. The gDNA present within the upper aqueous phase was precipitated using 1 ml 99 % ethanol. The resulting gDNA pellet was washed once with 70 % ice cold ethanol and then air dried. The pellet was resuspended in TE (10 mM Tris HCl, pH 8.0, 0.1 mM EDTA) buffer and stored at -80℃ till further use.

### Southern blot analysis

For Southern blot analysis about (5 -10) μg gDNA isolated from different *U. maydis* strains were digested with suitable combination of restriction enzymes. Following this the digested DNA were run on 0.8 % agarose gel at constant voltage (60 V) till the dye front reached the end of the gel. The gel was then washed once in water and incubated in 0.15 M HCl for 30 min under gentle rocking condition. Thereafter the gel was again washed in water followed by 30 min incubation in 0.5 M NaOH (pH ∼12.5). At this stage the pH of the gel was neutralized by incubating it in pH neutralization buffer (1 M Tris HCl pH 7.5, 1.5 M NaCl). Following a brief wash of the gel in water, the DNA molecules that were resolved on the gel were transferred on to the nylon membrane (Amersham Hybond N+, GE Healthcare) by capillary action, using the 20X-SSC buffer system (3 M Tris HCl pH 7.0, 341 mM trisodium citrate dehydrate). Following transfer, the DNA were crosslinked onto the membrane by immersing the membrane in 20X SSC buffer and exposing to UV in a UV cross linker. This was followed by incubation of the membrane in Southern hybridization buffer (0.5 M sodium phosphate buffer pH 7.0, 0.5 % skim milk, 7 % SDS) for 2 h at 65℃. The membrane was then incubated with digoxigenin (DIG) labelled suitable DNA probes at 65℃ overnight. Following probe binding the membrane was washed twice in Southern wash buffer (0.1 M sodium phosphate buffer pH 7.0, 1 % SDS) at 65℃ and once in DIG wash buffer (0.1 M malic acid, 0.15 M sodium chloride, 0.3 % Tween-20, pH 7.5) at room temperature. The membrane was then incubated in DIG II buffer (0.1 M malic acid, 0.15 M sodium chloride, 1 % skim milk pH 7.5) for 30 min at room temperature. At this stage the membrane was incubated with the anti-digoxigenin Fab fragments conjugated to alkaline phosphatase (Roche) at a dilution of 1:10000 in DIG II buffer for 30 min. After antibody binding the membrane was washed twice for 15 min each in DIG wash buffer and once for 5 min in DIG III buffer (0.1 M NaCl, 0.05 M MgCl_2_, 0.1 M Tris HCl, pH 9.5). Finally, the chemiluminiscence produced due to the activity of the alkaline phosphatase was detected by using CDP star (Roche) as the substrate.

### Plant infections

To observe the progress of infection maize plants of the variety Early Golden bantam (EGB) were infected with different *U. maydis* strains. For preparing the inoculum *U. maydis* strains were grown in YEPSL media at 28℃ till OD_600_ reached 0.8. The cells were then collected and resuspended in sterile water at a final OD_600_ 1.0. This cell suspension was then used to infect maize seedlings through syringe infection. The infected plants were kept at 24℃ in a temperature-controlled chamber with a 16 h day / 8 h night cycle and 70 % humidity for different periods of time.

### RNA isolation and quantitative reverse transcriptase polymerase chain reaction (RT-PCR)

RNA isolations from maize leaves and *U. maydis* strains were carried out using TRIzol reagent (Ambion, Life Technologies) following manufacturer’s protocol. Maize leaves either uninfected or infected with different *U. maydis* strains were collected and ground into fine powder using liquid nitrogen. To about 500 μl packed volume of ground tissue 500 μl TRIzol reagent was added and the suspension was incubated on ice for 10 min. In case of *U. maydis* cells, the sporidial pellets from about 4 ml axenic culture were lysed in 500 μl TRIzol using bead beating method. The TRIzol suspensions were then centrifuged at 16000 g for 10 min at 4℃. To the supernatant 100 μl chloroform was added and the resulting suspension was mixed vigourously. Following this the mixture was allowed to stand at room temperature for 2-3 min and then centrifuged at 16000 g for 10 min at 4℃. The RNA present in the upper aqueous phase was then precipitated using 250 μl isopropanol. The mixture was incubated at room temperature for 10 min and then centrifuged at 16000 g for 10 min at 4℃. The resulting RNA pellet was washed twice with 75 % ethanol, air dried and finally dissolved in nuclease free water. cDNA synthesis was carried out with either 1 μg DNaseI treated RNA from *U. maydis* axenic culture or 5 μg DNaseI treated RNA from maize tissue using Revert Aid reverse transcriptase (Thermo Scientific) following manufacturer’s protocol. Real time PCR analysis was done using TB Green Premix Ex Taq II (Tli RNase H Plus) from Takara in a CFX96 touch Real-Time PCR System (Bio-Rad) following manufacturer’s protocol. All cDNA samples used were diluted at least 20 times in nuclease-free water prior to PCR. Each real PCR reaction was carried out in two technical replicates and three biological replicates. The house-keeping gene, peptidyl prolyl isomerase (*ppi,* UMAG_03726) was used as the internal control.

### Visualisation of different developmental stages of U. maydis in planta

The progress of infection in maize plants infected with different strains of *U. maydis* was monitored by staining the infected tissues simultaneously with propidium iodide (PI) and wheat germ agglutinin conjugated to Alexa Fluor 488 (WGA-AF 488) as described in (Redkar et al., 2018). Briefly, the infected leaf tissues were excised from plants at different time points following *U. maydis* inoculation and treated with ethanol to remove chlorophyll. This was followed by a treatment with 10% KOH for 4 h that allowed softening of the leaf tissue. The tissues were then washed with 1X phosphate buffer saline (PBS) pH 7.4 and incubated with WGA-AF488 and PI in a vacuum infiltrator for 30 min with 5 min vacuum and 5 min rest cycles. WGA-AF488 stained fungal cell walls and PI-stained plant cell walls were visualized under a confocal microscope with proper illumination (WGA-AF488: Excitation/Emission at 495/519nm; PI: Excitation/Emission at 493/636 nm).

### Statistical methods

Statistical significance was computed in MS Excel. Significance between same group was assessed by paired two-tailed T-test, whereas significance between two different groups was assessed by unpaired two-tailed T-test. ns, *, **, ***, **** indicates P>0.05, p<0.05, p<0.01, p<0.001and p<0.0001 respectively.

### Purification of recombinant Hsp20

In order to carry out in vitro experiments demonstrating protein aggregation prevention activity associated with Hsp20, an N-terminal 6X His tagged version of the protein was expressed in *Escherichia coli* BL21 (DE3)-RIL harbouring the RIL^cam^ plasmid. Expression of recombinant His_6X_-Hsp20 was induced with 0.5 mM isopropylthio-β-galactoside (IPTG) at 37℃ for 3 h. Following induction, the cells were collected by centrifugation and resuspended in lysis buffer (50 mM Tris pH 8.0, 150 mM NaCl) supplemented with complete EDTA-free protease inhibitor cocktail (1 tablet/50 ml of lysate; Roche) and 1 mg/ml lysozyme (Sigma). Lysis was carried out by incubating the cell suspension on ice for 30 min followed by sonication using an ultrasonic processor (Cole Parmer, US). The lysed cells were then centrifuged at 24,303 g for 30 min (rotor SA-300, SorvallThermo) to remove cell debris. To purify recombinant His_6X_-Hsp20, the supernatant was applied to a Ni^2+^-NTA affinity column (Takara). Non-specifically bound proteins were removed by two successive wash of the column using lysis buffer supplemented with 10 mM and 20 mM imidazole respectively. The bound recombinant protein was eluted with lysis buffer containing 100 mM imidazole. The eluted fractions were monitored by running reducing SDS-PAGE and stored in aliquots at -80℃ till further use. For phase separation assay, recombinant Hsp20 was purified in phosphate buffer (50 mM sodium phosphate, 100 mM NaCl, pH 7.4).

### Protein aggregation protection assay

To assess any protein aggregation protection activity associated with Hsp20, in vitro protein aggregation protection assay was carried out using 10 μM lysozyme as substrate and 20 mM DTT as a chemical denaturant. Hsp20 was used in the reaction in 1:1 molar ration with lysozyme. Lysozyme incubated with DTT in the absence of Hsp20 served as the control sample. Protein aggregation in both the control and the test sample was monitored by measuring light scattering at 360 nm for 1 h at 25℃.

### Dynamic light scattering

To analyse the size distribution of Hsp20 at different temperatures or in the presence of SDS, dynamic light scattering (DLS) was performed using a Zetasizer Nano S (Malvern Instruments, Malvern, UK). The size distribution of 5 µM purified Hsp20 in 50 mM Tris (pH 8.0) and 150 mM NaCl buffer was measured at 25°C, 35°C, and 45°C. Additionally, size distribution measurements were taken both with and without 0.4 mM SDS.

The hydrodynamic radius (RH) was determined using the Stokes–Einstein equation:

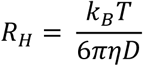

where k_B_ is Boltzmann’s constant, T is absolute temperature, η is the medium’s viscosity, and D is the translational diffusion coefficient of the particles. The data reported shows an average of 30 scans.

### CD spectroscopy

The far-ultraviolet (UV) circular dichroism (CD) spectrum of Hsp20 was measured at 25°C, 35°C, and 45°C, covering the range from 200 to 260 nm, using a quartz cell with a path length of 1 mm in a JASCO 150 CD-spectropolarimeter. Data were collected with a pitch of 1 nm and a scan rate of 100 nm/s. The far-UV spectrum was also recorded after treating Hsp20 with 0.4 mM SDS at 25°C. Each presented far-UV spectrum is the mean of three acquisitions. The spectra were deconvoluted using the BestSel software.

### Intrinsic tryptophan fluorescence measurement

Intrinsic tryptophan fluorescence of recombinant purified Hsp20 was measured at different temperatures (25°C, 35°C, and 45°C). The experiment was conducted using a JASCO F-8500 fluorimeter (JASCO International Co., Ltd., Japan), with samples excited at a wavelength of 280 nm. Fluorescence emission spectra were measured over the range of 300 nm to 500 nm, with data points collected at 1 nm increments. Additionally, tryptophan fluorescence was measured while decreasing the temperature and also in the presence, and absence of 0.4 mM SDS. This setup allowed for monitoring changes in tryptophan fluorescence, providing insights into the structural and conformational alterations of Hsp20 under different external conditions.

### Tagging of recombinant proteins

To conjugate Alexa-488, Alexa-532, and ATTO-655 fluorescent probes with purified recombinant proteins, a protocol following the manufacturer’s guidelines from Sigma-Aldrich (Saint Louis, MO, USA) was followed. Initially, a solution of purified recombinant proteins was prepared, and a sodium bicarbonate solution was added to create an optimal environment. The fluorescent probes’ stock solutions were prepared in anhydrous dimethyl sulfoxide (DMSO). The labeling process involved the slow addition of the fluorescent probes to the protein solution, followed by gentle stirring for 2 h. The separation of labeled protein from the unbound probe was achieved through gel filtration using a PD-10 column. The collected fractions then underwent a 24-hour dialysis against sodium phosphate buffer to remove excess probe and DMSO. The concentration and degree of labeling were assessed by measuring absorbance and fluorescence, with the Fluorophore-to-Protein (F/P) ratio indicating the efficiency of the labeling process. A final confirmation of successful protein labeling was obtained through SDS-PAGE, followed by scanning the gel using a Typhoon scanner (GE Healthcare, Danderyd, Sweden). This comprehensive protocol ensures efficient and optimized labeling of recombinant proteins, making them suitable for various assays and experiments requiring fluorescent detection.

### In vitro phase separation assay

The phase separation property of fluorescent labeled Hsp20 was assessed in the presence of varying concentration of PEG 8000 (2.5 %, 5 %, 10 % and 15 %). The concentration of Hsp20 was varied from 10 μM to 100 μM in 10 μM increaments. For the assay involving Hsp20 and actin together, 15 μM Alexa fluor 488 tagged Hsp20 and 5 μM atto 655 tagged actin were used. All the samples were incubated at 37℃ for 1 h and visualized under a confocal microsope.

### Fluorescence resonance energy transfer (FRET)

To assess the interaction between Hsp20 and actin, we employed Fluorescence Resonance Energy Transfer (FRET). Alexa 488-tagged actin at a concentration of 300 nM was titrated with Alexa 532-tagged Hsp20 at 25°C. The reaction mixture was excited at 490 nm, and emission spectra were recorded from 500 nm to 600 nm using a JASCO F-8500 spectrophotometer (JASCO International Co., Ltd., Japan). Emission spectra for both the donor and acceptor fluorophores were separated and analyzed using Igor Pro 6.0 software by WaveMetrics, USA. The decrease in donor fluorescence intensity was plotted against Hsp20 concentration, indicating the fraction of actin bound to Hsp20. Subsequently, this dataset was fitted to a mathematical equation for further analysis and interpretation.

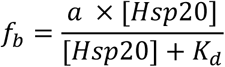

Where a is a constant, f_b_ is the bound fraction of Actin, and K_d_ is the dissociation constant. A control experiment was done where Alexa-488 tagged actin was titrated with Alexa-532 tagged lysozyme.

## Supplementary Information

Fig S1. Purification of His_6X_-Hsp20.

Fig S2. Titration of Alexa 488 tagged actin with Alexa 532 tagged Hsp20.

Fig S3. Hsp20 structure prediction using AlphaFold protein structure database.

Fig S4. Induced expression of b-gene rescues filamentation defect in AB33Δ*hsp20* mutant.

Fig S5. Subcellular localization of Hsp20 in *U. maydis*.

Video S1 & S2. Distribution of actin patches and cables in *U. maydis* SG200 wild type and *hsp20* deletion mutant.

Table S1. Strains used and generated in this study.

Table S2. Cloning strategies for different plasmid constructs used in this study.

Table S3. Primers used in this study.

## Acknowledgements

This study was funded by Science & Engineering Research Board/Anusandhan National Research Foundation, Govt. of India (Grant No. CRG/2021/000332) and Bose Institute intramural funding. A.M. and A.K. were supported by Department of Biotechnology, Govt. of India through DBT-JRF program.

## Author Contributions

A.G.^4^ and A.G.^5^ conceived the study. A.M., A.K. and K.B. performed most of the experiments with the help of A.R. and D.M. A.G.^4^ and A.G.^5^ analysed the data with the help of A.M. and K.B. A.G.^4^ and A.G.^5^ wrote the manuscript with inputs from A.M., K.B. and A.K. ^4^Anupama Ghosh, ^5^Abhrajyoti Ghosh

## Declaration of Interests

The authors declare no competing interests.

